# Off-resonance saturation as an MRI method to quantify ferritin-bound iron in the post-mortem brain

**DOI:** 10.1101/2021.03.22.436424

**Authors:** Lucia Bossoni, Ingrid Hegeman-Kleinn, Sjoerd G. van Duinen, Lena H. P. Vroegindeweij, Janneke G. Langendonk, Lydiane Hirschler, Andrew Webb, Louise van der Weerd

## Abstract

**Purpose:** To employ an Off-Resonance Saturation (ORS) method to measure the ferritin-bound iron pool, which is an endogenous contrast agent which can give information on cellular iron status.

**Methods:** An ORS acquisition protocol was implemented on a 7T preclinical scanner and the contrast maps were fitted to an established analytical model. The method was validated by correlation and Bland-Altman analysis on a ferritin-containing phantom. Ferritin-iron maps were obtained from post-mortem tissue of patients with neurological diseases characterized by brain iron accumulation, i. e. Alzheimer’s disease, Huntington’s disease and aceruloplasminemia, and validated with histology. Transverse relaxation rate and magnetic susceptibility values were also obtained for comparison.

**Results:** In post-mortem tissue, the ferritin-iron contrast strongly co-localizes with histological iron staining, in all the cases. Quantitative iron values obtained via the ORS method are in agreement with literature.

**Conclusions:** Off-resonance saturation is an effective way to detect iron in grey matter structures, while mitigating for the presence of myelin. If a reference region with little iron is available in the tissue, the method can produce quantitative iron maps. This method is applicable in the study of brain diseases characterized by brain iron accumulation and complement existing iron-sensitive parametric methods.

## Introduction

Physiological iron is distributed heterogeneously within the brain, with the structures richest in iron being localized within the basal ganglia^1^. The distribution of iron throughout the brain is likely related to its fundamental biological processes, which involve oxygen transport, DNA synthesis, and mitochondrial respiration^2^. In contrast, the dysregulation of iron homeostasis is associated with neurotoxicity via the formation of reactive oxygen species (ROS)^2–6^, lipid peroxidation^7^, and ferroptosis^8^, a form of cell death characterized by phospholipid oxidation^9^. Iron is also thought to trigger inflammation which can, in turn, lead to additional cellular iron release^10^.

Brain iron accumulation is a common phenomenon of several neurodegenerative diseases, despite their different pathological hallmarks. Examples are Alzheimer’s disease^2,11^, Huntington’s disease^2,4,12^, and the neurodegeneration with brain iron accumulation (NBIA) group of disorders of which aceruloplasminemia (ACP) is a rare phenotype^13^.

One of the primary cytoplasmic proteins responsible for intra- and extracellular iron storage is ferritin^14^. Ferritin is composed of a 24-subunit shell (apoferritin) which plays three important roles in iron regulation: it binds cytoplasmic Fe^2+^; it oxidizes Fe^2+^ into Fe^3+^ (as antioxidant protection); and it stores iron in the form of a biocompatible nano-mineral^15^, ferrihydrite, or a mixture of ferrihydrite and other minerals^16,17^. Since the core of ferritin can bind up to ~5000 iron atoms, the nanoparticle’s saturated magnetic moment can reach ~300 μ_B_^18^. Ferritin-bound iron levels constitute ~ 80% of total non-heme iron^19^ and therefore ferritin is commonly used as a reporter of tissue iron status. Although ferritin-bound iron is unlikely to contribute directly to oxidative stress, its levels are related to the labile iron pool^20,21^, which is implicated in neurotoxicity.

MRI is very sensitive to the magnetic field perturbation caused by iron ions and iron nanoparticles which have superparamagnetic properties. In the presence of iron, diffusing water molecules probe a broad distribution of magnetic fields. Consequently, a time-dependent phase is accumulated, leading to reduced T_2_ and T_2_^*^ relaxation times^22–24^. Ferritin-bound iron is the (physiological) iron form having the largest effect on the MRI signal^19^ due to its large susceptibility and abundance. The most sensitive MR methods for the detection of iron are the Field Dependent Transverse Relaxation Rate Increase (FDRI)^25–27^, R_2_^*^ and R_2_ mapping^28,29^, and Quantitative Susceptibility Mapping (QSM)^30^. Although quantitative maps of transverse relaxation rates and tissue magnetic susceptibility have already shown benefits in the clinic^22,31^, these parameter are also influenced by myelin content^32–34^, fiber orientation^35^, tissue microstructure^36^, neuronal loss^29^ and iron aggregation^37,38^. Additionally, QSM reconstructions can be affected by the non-local nature of the inverse problem that relates magnetic field perturbation to magnetic susceptibility, and this impairs quantitative analysis.

In this work, we show how an off-resonance saturation (ORS) method, earlier introduced for the detection of iron-oxide contrast agents^39,40^, can be employed to assess ferritin-bound iron levels in the post-mortem brain and overcome some of the limitations of conventional mapping methods.

We validated the method in a brain phantom, and we employed it to characterize the ferritin-bound iron levels in post-mortem material obtained from three patients affected by neurological diseases associated with increased brain iron. We assessed the ferritin-bound iron maps against QSM and R_2_^*^ maps, and histopathological staining for ferric iron, ferritin and myelin.

## Methods

### Tissue selection and phantom preparation

A ferritin-agar phantom was prepared by adding horse spleen ferritin (Sigma, ref. F4503) to a 1.5% agar solution. Phantoms with different iron concentrations were prepared: 0 mM, 1.3 mM, 4.0 mM, 7.0 mM, 10.3 mM and 15.1 mM. The iron concentration was estimated by multiplying the ferritin concentration by the average loading of the protein i.e. 2500 iron/ferritin^41^.

Formalin-fixed tissue blocks extracted from three diseased brains were studied. Brain material from a patient with Huntington’s disease (HD), a patient with Alzheimer’s disease (AD) and a patient with aceruloplasminemia (ACP) was obtained from the Pathology department of the LUMC, the Netherlands Brain Bank, and the Erasmus Medical Center, respectively. Patient’s informed consent was obtained from each respective brain bank. For each case, a disease-relevant area was chosen: the striatum for the HD case; the striatum and the globus pallidus for the ACP case; and the middle temporal gyrus for the AD case. The tissue blocks were rehydrated in phosphate buffered solution (PBS) for 24 hours prior to the scan, to partially restore transverse relaxation times. Subsequently, the tissue was scanned in a hydrogen-free solution (Fomblin^®^ LC08, Solvay).

### MRI data acquisition

MRI data were acquired on a 7T preclinical scanner (Bruker Biospin, Ettlingen, Germany) using a 38 mm linear birdcage transmit-receive coil (T10327V3) with eight legs.

To compare the iron maps obtained with ORS to standard QSM and R_2_^*^ maps, R_2_^*^-weighted images were acquired using a 3D multiple gradient echo (MGE) sequence. The echo time (TE) was optimized for the tissue block under study since the transverse relaxation (and iron load) largely varies per region and disease. The acquisition parameters are summarized in Table 1.

**Table 1.** Summary of the acquisition parameters for the MGE sequence, with eight echoes (N_echo_). TE_1_ refers to the first echo time, ΔTE refers to the inter-echo time. TR is the recovery time and FA the flip angle. Abbreviations: HD: Huntington’s disease, ACP: aceruloplasminemia, AD: Alzheimer’s disease.

| Sample | Pixel size (mm<sup>3</sup>) | FA (degrees) | Averages | TR (ms) | TE<sub>1</sub> (ms) | N<sub>echo</sub> | $\Delta TE$ | Total scan time |
| --- | --- | --- | --- | --- | --- | --- | --- | --- |
| HD | (0.15)<sup>3</sup> | 25 | 20 | 107.3 | 5.2 | 8 | 4.34 | 3h26m |
| ACP | (0.15)<sup>3</sup> | 25 | 20 | 130.4 | 1.96 | 8 | 2.21 | 3h55m |
| AD | (0.15)<sup>3</sup> | 25 | 20 | 107.3 | 5.2 | 8 | 4.34 | 3h26m |

ORS images were derived from 2D steady state free precession (SSFP) images (FISP-FID) with TE/TR=3/6ms; inter-scan delay TR =2 s; FA= 10 degrees. A ‘control’ magnitude image (M_control_), without saturation pulses, was also acquired. Saturated images (M_sat_) were produced by a saturation module consisting of six hyperbolic secant pulses of bandwidth 500 Hz (B_1_=1.36 μT), followed by a spoiler gradient of 3 ms duration. The off-resonance saturation pulses were phase-cycled to improve the spoiling. A list of Msat was obtained upon varying the frequency of the off-resonance pulse. Details of the off-resonance acquisition are reported in Table 2.

**Table 2.** Summary of the acquisition parameters for the ORS-SSFP acquisition. Symbols: BW: ORS Pulse bandwidth; ν0: frequency of the off-resonance saturation pulse; Δν0:interval between ORS frequencies, HD: Huntington’s disease, ACP: aceruloplasminemia, AD: Alzheimer’s disease.

| Sample | BW [Hz] | $\nu_0$ range [Hz] | $\Delta\nu_0$ [Hz] | Total scan time | Pixel size (mm<sup>3</sup>) | Reference for contrast calculation (ROR) |
| --- | --- | --- | --- | --- | --- | --- |
| Phantom | 500 | 250-649 | 7 | 1.3h | 0.3x0.3x2 | Agar-only compartment |
| HD | 500 | 250-700 | 5 | ~8h10min | 0.15x0.15x2 | Internal capsule |
| ACP | 500 | 250-700 | 5 | ~8h10min | 0.15x0.15x2 | Internal Capsule |
| AD | 500 | 250-700 | 5 | ~8h10min | 0.15x0.15x2 | Cortical white matter |

Raw data acquired in this study are freely available at the link: http://dx.doi.org/10.17632/nkznjjnwpw.1

### MRI Data Processing

MGE-magnitude images were fitted on a pixel-by-pixel basis to a mono-exponential decay function^23^, after ruling out the presence of multi-components, using the Levenberg–Marquardt curve-fitting algorithm to derive the transverse relaxation rate R_2_^*^. The noise floor was reached in the last 2 or so echoes in some regions of the tissue (ACP) containing extremely high iron accumulation but did not affect the overall fitting quality. To more accurately fit the data, we excluded the pixels with intensities at the noise level.

Susceptibility values (χ) were obtained from the phase images with the STI-Suite software (version 2.2). Phase unwrapping of the measured phase images and removal of the background field were done with iHARPERELLA^42^, while magnetic susceptibility estimation and streaking artifact correction were performed with the ‘iLSQR’ algorithm^43^. Raw susceptibility values are reported, as no reference region was available in these small tissue samples.

Figure 1 illustrates the pipeline of the ORS analysis. ORS images (M_ors_) were obtained from the subtraction: M_ors_ = M_control_ - M_sat_. Contrast (C) maps were obtained by subtracting the mean intensity of M_ors_ in a region of reference (ROR) from the Mors40. Under the assumption that the ROR contains no iron, the contrast (C) can be written as the product of the fraction of protons (φ) saturated by the ORS pulses and the intensity of the control image (C=M_control_ φ). In our work, only positive frequency offsets were chosen, to exclude Nuclear Overhauser Effects (NOE)-related artifacts^44^ and to speed up the acquisition. The saturated fraction of protons, as derived by Delangre *et al.*^40^, is given by:

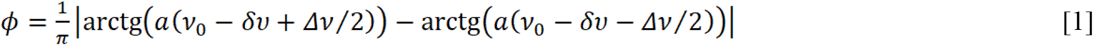

**Figure 1.**
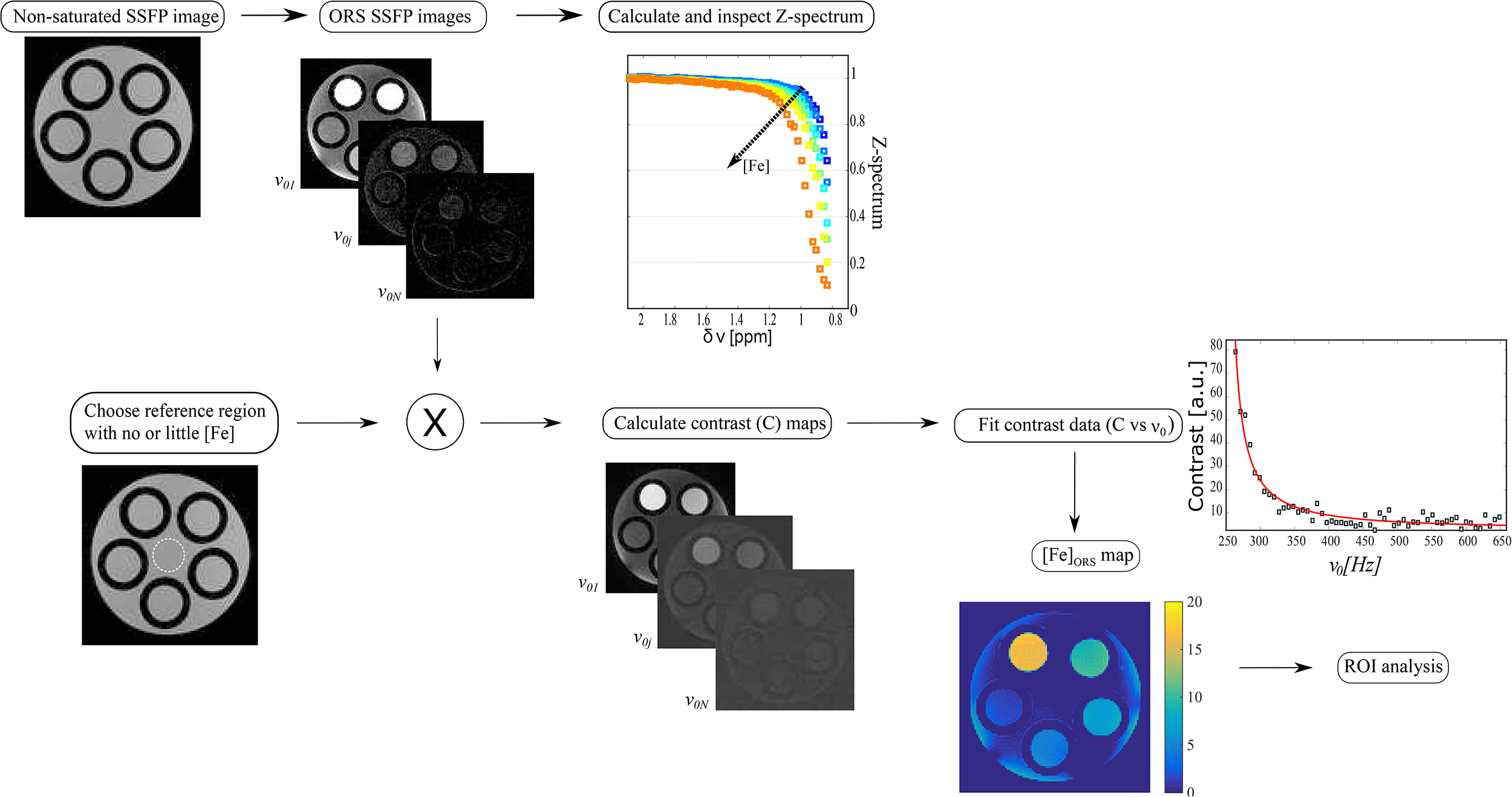
Analysis pipeline. After the acquisition of the non-saturated images (‘control’), the saturated images are acquired, and the ORS image is derived. The Z-spectrum is shown, but was not used in the analysis. The region of reference (ROR) is shown here as a white dotted line in the bottom row. The contrast maps are obtained from the ORS image and the mean intensity in the ROR, and fitted to the model discussed below. An example of the fit is shown in the right panel. The fit is the red solid line overlapped on to the experimental data (blue squares). Finally, a map of iron concentration in mM is obtained ([Fe]_ORS_ map).

Where 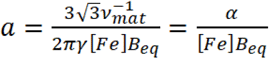, γ is expressed in MHz/T, 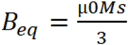 is the stray field (in Tesla) at the equator of a nanoparticle with saturation magnetization *M*_*s*_, [Fe] is the iron concentration (in mM), and *v*_*mat*_ is the molar volume of ferrihydrite^45^ and equal to 5.4×10^−6^ m^3^/mol=5.4×10^−6^ mM^−1^, and α=3.6 x 10^−3^ T (Hz mM)^−1^ (See Supplementary material for further details). A frequency offset (*δv*) was added to account for resonance frequency shifts. Since N=6 saturation pulses were used to enhance the saturation^46^, the final expression for the contrast becomes:

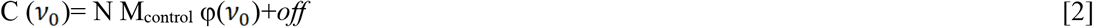

M_control_ is weighted by the proton density, relaxation times, and flip angle^47^. The offset (*off*) term in the equation above was introduced to account for residual MT effects and imperfect saturation. Finally, the contrast map was fitted to equation [2] to derive the iron concentration, with the equatorial field as a fixed parameter. The agreement between the fitted iron concentration and the known iron concentration was assessed with correlative and Bland-Altman analyses.

### Histopathological validation

The same tissue blocks used for MRI were employed for histology. Tissue blocks were embedded in paraffin and sliced with a microtome into 5 μm and 20 μm thick sections. The 20 μm sections were employed for non-heme iron detection (Meguro staining), according to Van Duijn *et al.*^48^. After deparaffinization, the tissue sections were incubated for 80 min in 1% potassium ferro-cyanide, washed, followed by 100 min incubation in methanol with 0.01M NaN_3_ and 0.3% H_2_O_2_. Subsequently, sections were washed with 0.1M phosphate buffer followed by 80 min incubation in a solution containing 0.025% 3’3-diaminobenzidine- tetrahydrochloride (DAB, Sigma-Aldrich) and 0.005% H_2_O_2_ in 0.1M phosphate buffer. The reaction was stopped by washing with tap water. The 5 μm slices were used for additional staining: non-heme (mostly trivalent) iron was detected with Perl’s Prussian blue (Merck 1.04984.0100); and immunohistochemical detection of myelin was done with anti-myelin PLP antibody (Bio-Rad, MCA 839G) with second antibody Rb-aMs/biotin (DAKO) for 1h at room temperature, followed by ABC (avidin-biotin-complex/HRP, ABC Elite Kit, Vector) for 30 min at room temperature. Immunohistochemical detection of ferritin was done with ferritin antibody (Bethyl A80-140, dilution 1:1000, overnight incubation at room temperature), with as second step antigoat/biotin (Betyhl A50-204B, dilution:1:1000, 1h incubation at room temperature), followed by ABC incubation for 30 min at room temperature. After the ABC treatment, the tissue was rinsed three times with phosphate buffered saline and incubated in 0.05% DAB (Sigma-Aldrich) with 15 μl 30%H_2_O_2_/100ml for 5-10 minutes. After rinsing several times with demineralized water, the slices were counterstained for 30 s with Harris haematoxilin and washed for 5-10 mins with tap water. Finally, the tissue sections were dehydrated with ethanol 70%, 96%,100%, and xylene.

### Correlation between histology and ORS imaging

To quantify the degree of agreement between the histological staining for iron with the ferritin-bound iron concentration derived from the ORS method, oval regions of interest (ROI, N=30) were drawn in the grey matter regions of the ferritin-bound iron maps and ORS maps. The iron staining image was manually co-registered (affine transformation) to the MRI maps with the TrackEM2 plugin in ImageJ. The registered image stack was converted to an 8-bit greyscale image. Oval ROIs were carefully drawn and propagated between the iron histological map and the MRI map with the ROI manager tool. Mean grey values from the predefined ROIs were quantified^32^, after inspecting for the precise placement of the ROIs. Pearson’s correlation coefficient (ρ) of the association between histological staining intensity and the ferritin-bound iron map was calculated per case.

## Results

### Validation on ferritin phantom

Figure 2 shows the iron map obtained for the ferritin-loaded phantom and the ROI analysis on the sample’s compartments. The region of reference (ROR) was drawn in the middle of the phantom containing only agar (Figure 1). The best-fitting results were obtained when fixing B_eq_ = 1 mT, which is very close to the theoretical value for ferritin (see Supplementary Information). The fitted iron value agrees well with the nominal iron concentration, as assessed by the correlative plot in Figure 2C. Furthermore, the agreement between the iron concentration derived from the ORS map and the nominal iron concentrations was assessed by Bland-Altman analysis (Figure 2D)^49,50^, considering the latter parameter as the reference value. No data fell outside of the limits of agreement and no systematic bias was detected, at the 5% level of significance.

**Figure 2.**
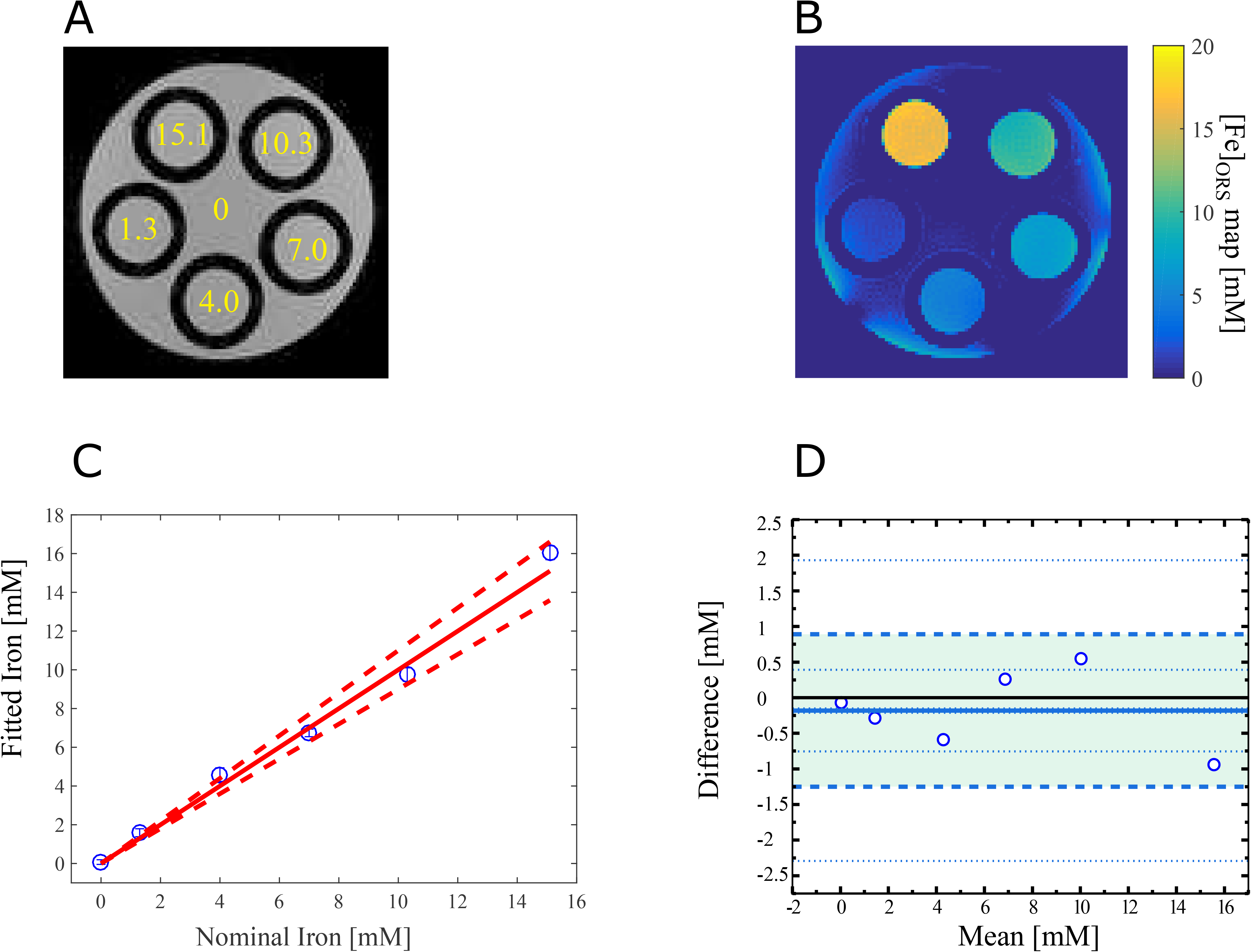
Validation on a ferritin phantom. Panel A shows the control image with the nominal iron concentration in mM, per compartment. Panel B shows the ferritin-bound iron map in mM, as obtained with the ORS sequence and the fitting method discussed above. The bright outer rim in the iron map is likely caused by B1 inhomogeneities, since a linear eight-leg birdcage coil with a high filling factor was employed. Panel C shows a comparison between the nominal iron concentration and the fitted one from the method discussed in this work. Blue data with error bars represent the mean and standard deviation of the iron concentration in the ROI drawn in each sample compartment. The red solid line is the identity line and the dashed lines account for the spread in iron loading as reported in literature for horse spleen ferritin from the same producer. Panel D is the Bland-Altman plot of the same data. The black-solid line is the line of equality. The blue line is the bias. The dashed lines enclosing the green area are the limits of agreement. The dotted lines mark the 95% confidence intervals for each estimated statistical quantity.

### Application to post-mortem material

Figure 3 shows the results obtained for the middle temporal gyrus of the AD case. A comparison is shown between the MRI parametric maps, i.e. R_2_^*^, QSM and the ORS-based ferritin-bound iron map (termed [Fe]_ORS_ map in the rest of the manuscript) and histology, i.e. myelin, iron and ferritin. Elevated R_2_^*^ levels are seen in the myelin-rich white matter and in the more superficial cortical layers. In the QSM map, negative susceptibility is predominantly detected in the white matter. The [Fe]_ORS_ map shows a strong localization of the iron in the cortex and (partially) in the white matter, thus largely mirroring the QSM paramagnetic contrast and the histological staining of iron and ferritin (protein), while it appears poorly correlated to the myelin histological staining. Across all tissue blocks, the Perl’s staining was considerably weaker than then Meguro staining, as earlier reported^48,51^, and correlated less with the ferritin map, therefore our quantitative analysis was based on the Meguro staining for iron, only.

**Figure 3.**
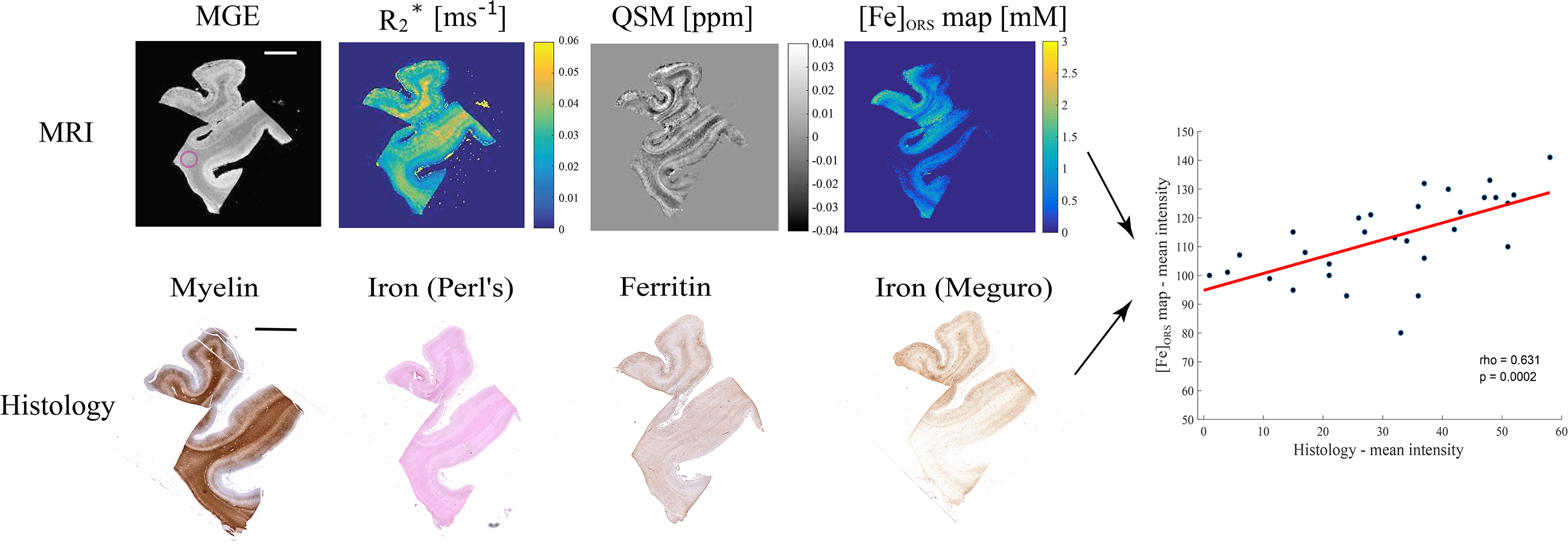
Comparison of the quantitative MRI and histological methods to assess tissue iron load on the Alzheimer tissue block: Top row: MRI R_2_^*^, QSM and ferritin-bound iron mapping. Bottom row: histological staining for myelin, iron (Perl’s and Meguro) and ferritin, Scalebar: 5 mm. The 4^th^ echo of the MGE image is shown for anatomical reference. The pink circle in the MGE image encloses the region of reference (ROR). The panel on the right shows the result of the linear regression analysis between the ROI mean intensity of [Fe]_ORS_ map and the histological staining for iron. The Pearson’s correlation coefficient and p-value are reported on the graph.

The tissue block containing the striatum of the HD case (Figure 4) shows high R_2_^*^ primarily in the putamen (PU), some connecting fibers of the internal capsule (IC), and a thin stripe of the atrophied caudate nucleus (CN). The susceptibility map looks heterogeneous, but with an overall positive susceptibility in the PU and CN. The IC shows a characteristic striped appearance of alternating positive and negative susceptibility. The [Fe]_ORS_ map shows a localized increase of iron in the PU, a thin stripe of the CN and of the external capsule. This map mirrors the iron staining, with the exception of the external capsule and some connecting fibers of the internal capsule, which show no contrast in the [Fe]_ORS_ map, probably due to partial volume effects. The histological staining for ferritin, iron and the [Fe]_ORS_ map are largely opposite to the myelin-positive regions, which display negative susceptibility in the QSM map. Note that the [Fe]_ORS_ map shows iron contrast in a substantial part of the external capsule, possibly due to iron in myelin.

**Figure 4.**
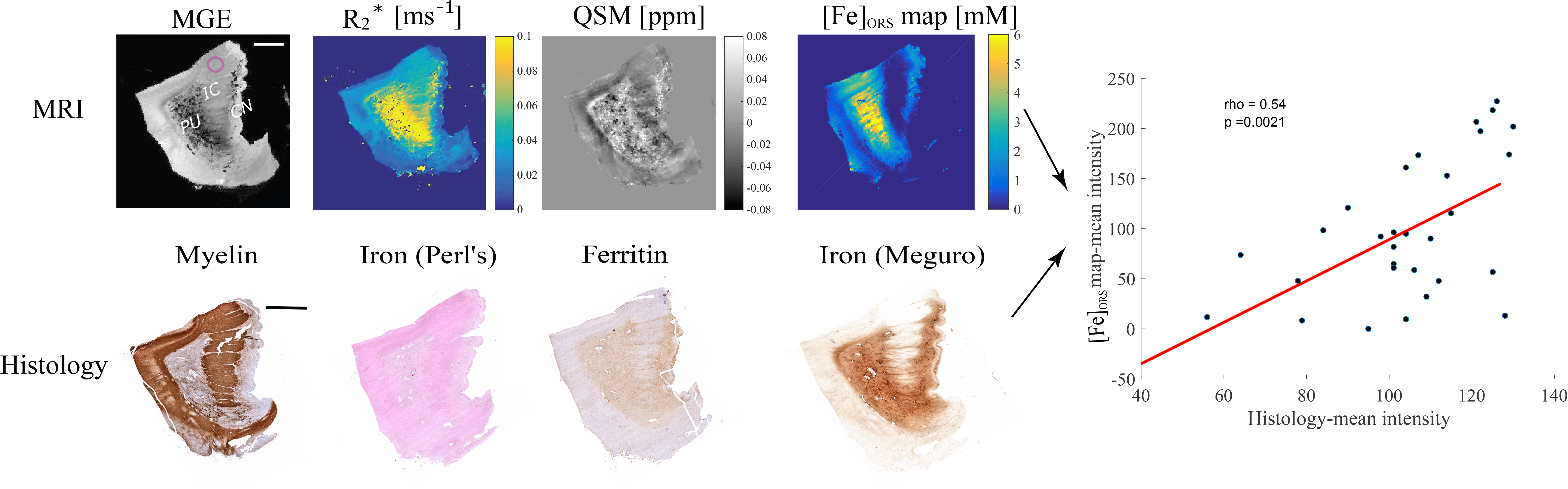
Comparison of the quantitative MRI and histological methods to assess tissue iron load in the Huntington’s diseases tissue block (labels as above): Scalebar: 5 mm. Abbreviations: IC= internal capsule, PU=putamen, CN=caudate nucleus. The MGE image displays the third echo.

Finally, the tissue block obtained from the ACP patient (Figure 5) shows high R_2_^*^ in the putamen, some connecting fibers of the internal capsule, the caudate nucleus and the globus pallidus (R_2_^*^~0.3-0.6 ms^−1^), similarly to the HD case. The QSM map is relatively smooth in the white matter regions, while the grey matter is heterogeneous and presents a ‘patchy’ appearance (also observed in the MGE images) which hinders quantitative analysis. The [Fe]_ORS_ map on this tissue block shows diffuse iron across both white and grey matter structures, except for the superior part of the internal capsule, where the ROR was drawn. The histological iron staining (Meguro) appears very intense across the whole slice, with a slightly higher intensity in the caudate nucleus and the globus pallidus, which is also captured by the [Fe]_ORS_ map. In contrast to the previous cases, the ferritin staining is rather weak across the whole slice, as discussed below. Additional histological results on all the tissue blocks are found in the Supplementary material (Figures S2-S5).

**Figure 5.**
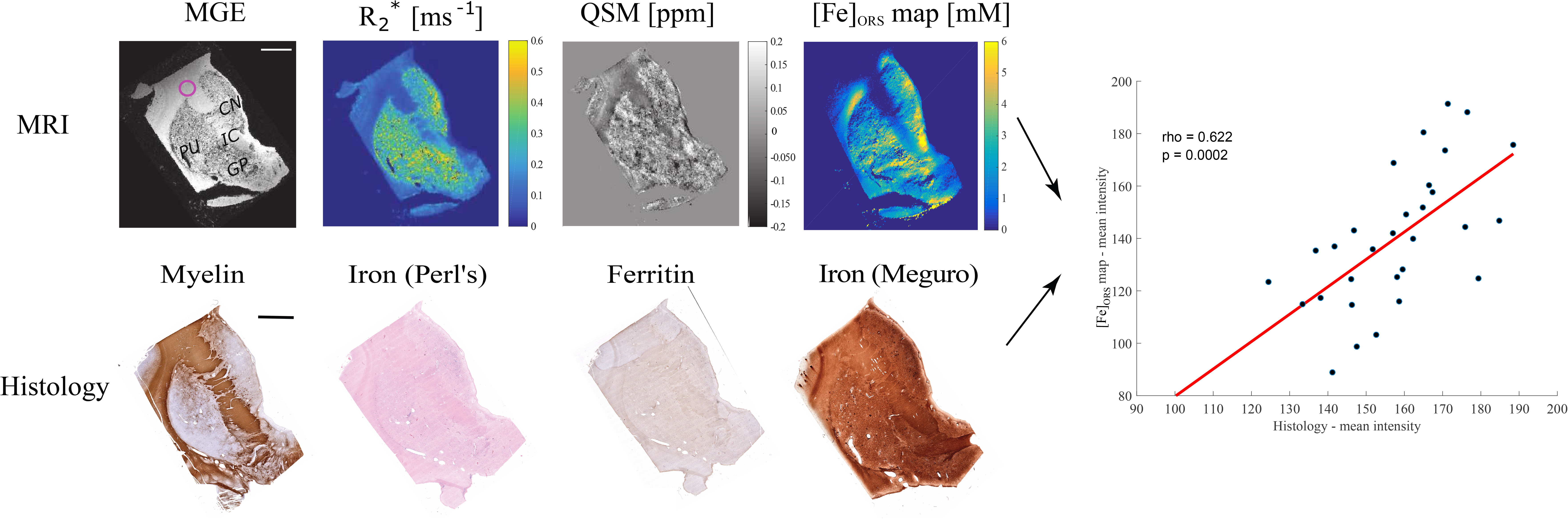
Comparison of the quantitative MRI and histological methods to assess tissue iron load on the aceruloplasminemia tissue block (labels as above): Scalebar: 5 mm. Abbreviations: IC= internal capsule, PU=putamen, CN=caudate nucleus, GP=globus pallidus. The MGE image displays the first echo.

## Discussion

In this work we demonstrate how a method based on diffusion-mediated off-resonance saturation (ORS)^39^ can detect ferritin-bound iron concentrations from post-mortem brain material of an Alzheimer’s (AD), Huntington’s (HD) and aceruloplasminemia (ACP) patient. This method can be translated in vivo to evaluate tissue iron load and therapeutic efficacy on a pathophysiological level.

Water molecules diffusing around iron particles can be selectively excited by ORS pulses, based on the correspondence between frequency offset and distance from the nanoparticle: the larger the frequency offset, the closer the protons are to the dipolar-field source. It has been shown that the positive contrast obtained by this method is distinct from, and can be observed also in the presence of, MT effects^39^. Although the ORS-method was originally introduced to quantify/detect superparamagnetic iron-oxide nanoparticles (SPION), here we show that ORS can also be employed to assess the presence and the concentration of ferritin-bound iron, by virtue of the super-paramagnetic properties of the protein and its high concentration across the brain. Our results on the agarose phantom show good agreement between nominal and fitted iron concentrations and support the use of the method for iron quantification.

Visual comparison between the parametric MRI maps acquired in this study shows some advantages of the ORS method: i) iron load appears highly localized in specific brain regions (mainly the gray matter); ii) the white matter structures, which are highly myelinated, are predominantly masked out in the [Fe]_ORS_ maps, except for the ACP tissue where iron appeares largely diffuse throughout the whole slice; iii) the [Fe]_ORS_ map largely mirrors the QSM map, when the latter is free from artefacts; iv) the strength of the association between the [Fe]_ORS_ maps and the iron staining was moderately high in all cases (ρ>0.5), and independent on the disease type.

The appearance of the ferritin-bound iron map in the tissue block obtained from the AD patient suggests that, when looking at the temporal lobe, iron preferentially accumulates in the cortical grey matter. This is in agreement with previous R_2_^*52,53^ and QSM studies^54^ reporting increased cortical iron levels in patients with AD or mild cognitive impairment, which was associated with AD pathological hallmarks, and an increased risk of cognitive decline. Magnetometry studies carried out on both sporadic and genetic types of AD have also found that ferritin-bound iron is more abundant in the AD group than in age- and gender-matched healthy controls^55,56^.

Previous analytical studies have detected significant increase in the *total* iron content of the temporal lobe of AD patients with respect to controls^57,58^. However, the absolute iron concentrations ranged from ~30 ng/mg^58^ to 120 ng/mg (wet weight)^59^, since age, gender, disease state and technique sensitivity can impact the measured iron values. The highest cortical ferritin-bound iron concentrations detected in our tissue block were 1.37 ± 0.25 mM (mean ± sd), corresponding to 76.45 ± 13.95 ng/mg, which is in agreement with the estimate of ferritin-bound iron by magnetometry in the AD temporal cortex^56^, after correcting for dry weight mass-loss.

The ferritin-bound iron map of the tissue block from the HD patient shows an increase of iron in the putamen and part of the atrophic caudate nucleus. Alterations in brain iron metabolism with increased iron accumulation have been previously identified by MRI in the striatal nuclei of patients with Huntington’s disease^26,60^. Iron accumulation in these structures seems to occur early on during the disease process^61^, suggesting a key role of iron in the initiation and progression of the disease^11^. In fact, in vivo MRI studies at both 3T and 7T have shown that iron accumulation^62^ is associated with tissue atrophy of basal ganglia structures and is already present in the premanifest phase of HD^61,63,64^. Our mean fitted iron concentration in the putamen was 4.66 ± 1.35 mM, equivalent to 260.0 ± 75.33 ng/mg, which agrees well with iron concentrations measured by inductively coupled mass spectrometry^12^.

Finally, the ferritin-bound iron map of the ACP tissue block shows a very diffuse iron load, with mean iron values of 2.57 ± 1.45 mM, in the putamen, which is about 50% lower than our recent magnetometry study which quantified ferritin-iron in the same structure of the same patient^65^. This is likely due to the combination of very fast relaxation times and the lack of an appropriate ROR in the slice. In contrast to AD and HD, iron accumulation in ACP is directly related to its genetic background, and results from the absence of functioning ceruloplasmin^13^. The lack of ceruloplasmin-mediated oxidation of ferrous iron (Fe^2+^) to ferric iron (Fe^3+^) impairs iron efflux from astrocytes, and leads to massive iron accumulation within these cells, while neurons that are mainly dependent on the supply of iron from astrocytes are probably iron-starved^13,66^. Although ferritin-bound iron appears to be by far the most abundant iron form in the aceruloplasminemia brain^65,67,68^, these observations are based on patients with end-stage aceruloplasminemia, and it remains unclear how the increase in either total iron levels, as suggested by previous R_2_^* 13,69,70^, and QSM studies^69^, or in the amount of ferritin-bound iron, are associated with the clinical course of the disease. The apparent contradiction with the weak ferritin staining for the tissue block can perhaps be resolved by considering that the long formalin fixation (~ 5 years) can impact the tissue quality and staining efficiency.

Different MRI methods, in addition to those already mentioned, have been used to estimate the effect of ferritin nanoparticles on the MRI parameters. In fact, ferritin-bound iron has long been identified as the main source of iron-driving contrast, given the large abundance of the protein in the brain and its magnetic properties. The magnetic susceptibility of a single ferritin protein was estimated as χ = 520 ppm for a fully loaded particle^19^, although this value might be an upper limit^18,71^.

Ferritin-iron displays a peculiar linear inverse dependence of T_2_ with B_0_ field^72^, a trend that is retained in brain tissue and is attributed to the ‘fingerprint’ of iron stores. This characteristic is exploited by the Field Dependent Transverse Relaxation Rate Increase (FDRI) metrics^73^.

A method based on direct saturation to sensitize image contrast based on iron load^74^ has demonstrated a linear relation with the tissue iron content and an improved gray matter-white matter contrast with respect to T2-weighted images. The ORS method here proposed differs in the acquisition protocol, with a much larger ORS frequency range being probed here, and in the data post-processing.

When inspecting the ferritin-bound iron maps, some caveats should be considered. Firstly, our method assumes that the only source of contrast are ferritin nanoparticles carrying ferrihydrite in the core. This is a simplification, as ferrihydrite is probably also included in hemosiderin^75^. Also, magnetite/maghemite is located in the brain either outside or within ferritin^16,76^. Recent studies, though, have shown that these additional minerals are approximately three orders of magnitude less abundant than ferritin-bound iron.^55,56^

Secondly, we assumed that the iron loading of each ferritin protein was approximately equal to half of the maximum filling capacity of the homonymous protein, while lower iron loading ranges, i. e. between 1500-1850 iron ions within each ferritin protein, have been reported for AD brain tissue and controls^77^. However, since the magnetization of the ferritin particle would not, or only minimally, depend on the iron loading, this does not significantly affect the total ferritin-iron concentrations reported here.

Additionally, the ferritin-bound iron maps were obtained from contrast maps that were referenced to a region (ideally) without iron. Therefore, the iron concentrations here displayed cannot be considered as absolute. This is especially clear in the case of the ACP tissue block, where the ferritin-bound iron map shows comparable iron concentrations to the HD case, despite the striking difference in the Meguro staining. This is probably due to the lack of a good ROR in the ACP tissue block.

Finally, T_2_ maps (not acquired in this study) can provide additional information on the degree of iron accumulation and, in combination with the T_2_^*^ maps, could offer valuable information on the effective size of iron-rich compartments^78^.

In conclusion, we adapted an ORS method^46,39^ to quantify the ferritin-bound iron pool in the post-mortem brain tissue of three patients affected by neurological diseases associated with increased brain iron. This method can aid the interpretation of R_2_^*^ and QSM maps, especially when these are confounded by reconstruction artifacts or the co-presence of iron and myelin. The accuracy of the ferritin-bound iron map depends on the availability, within the tissue, of a region without (or with little) iron content with respect to the region of interest. We foresee that this method will find use in the study of the progression of neurodegenerative diseases characterized by brain iron accumulation and the assessment of iron chelation therapy.

## Author contributions

**Lucia Bossoni:**Conceptualization, Methodology, Software, Investigation, Writing – Original Draft, Visualization, Project Administration; **Ingrid Hegemann-Kleinn**: histological analysis; **Sjoerd van Duinen**: supervision of the histological analysis; **Lena H. P. Vroegindeweij**: Resources, Writing – Review & Editing; **Janneke G. Langendonk**: Resources, Writing – Review & Editing; **Lydiane Hirschler**: Software, Writing – Review & Editing; **Andrew G. Webb:**Methodology, Resources, Writing – Review & Editing; **Louise van der Weerd:**Methodology, Resources, Writing – Review & Editing.

## Acknowledgements

The authors are grateful to E. Suidgeest, E. Ercan and M. Bulk for assistance in the lab and useful discussions.

## Funding

This study was supported by the Netherlands Organization for Scientific Research (NWO) through a VENI fellowship to L.B. (0.16.Veni.188.040).

## Declarations of interest

none.

## Notes

### Competing Interest Statement

The authors have declared no competing interest.

