## Supplementary Materials for "Off-resonance saturation as an MRI method to quantify ferritin-bound iron in the post-mortem brain"

### Calculation of the parameters used in equation [1] of the main manuscript.

The molar volume for ferrihydrite was obtained from the following equation:

$$\nu_{mat} = \frac{M_{mol}}{8\rho}$$

where $M_{mol} = 0.17\ kg/mol$ is the molar mass of ferrihydrite and $\rho = 3960$ kg/m$^3$ is its mass density. The numerical factor at the denominator accounts for the number of iron atoms in the formula unit of the core mineral $(FeOOH)_8$[1].

To estimate the equatorial field of ferritin we first started from the theoretical value of $\chi$=520ppm [1]. Therefore

$$B_{eq}[T] = \frac{\mu 0 M}{3} = \frac{\chi B}{3}$$

leading to ~1.3 mT. A slightly lower result (1 mT) was obtained from magnetometry data: M(7 T, 297 K) ~0.6Am$^2$/kg$^2$. We used this second value in our analysis.

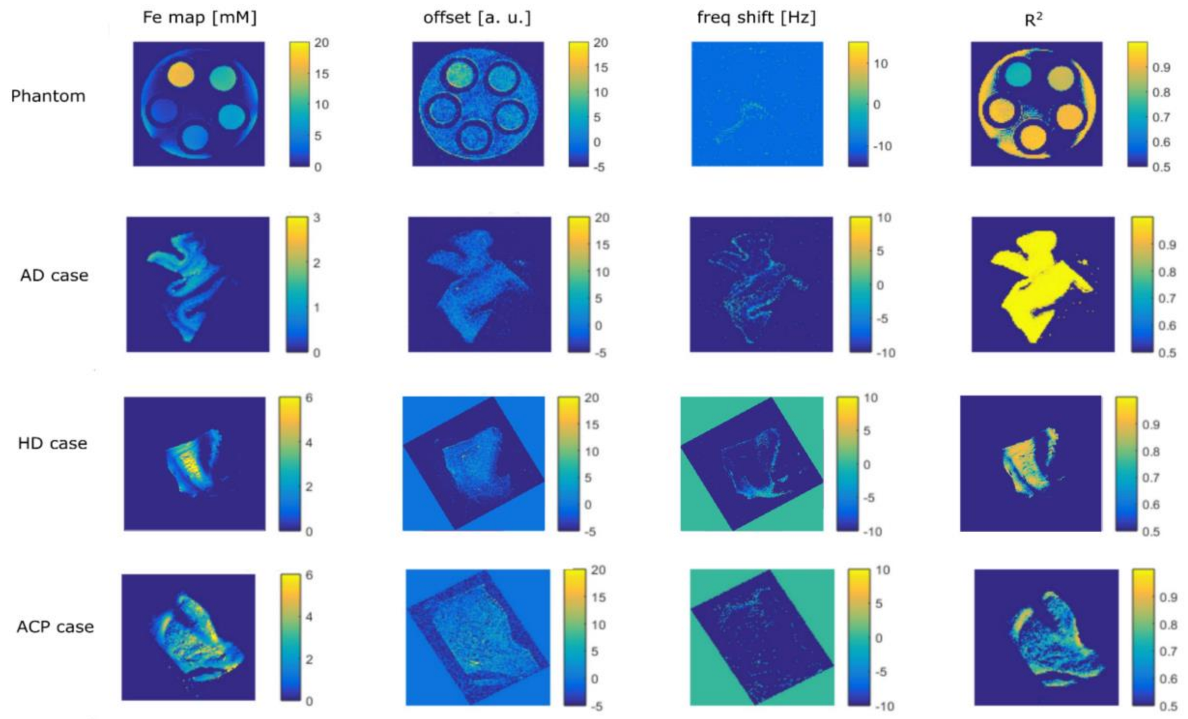


***Figure S1***. *Illustration of the quality of the fit and fitting parameters for the three tissue blocks and the phantom. See equation [2] of the main manuscript for the description of the parameters.*

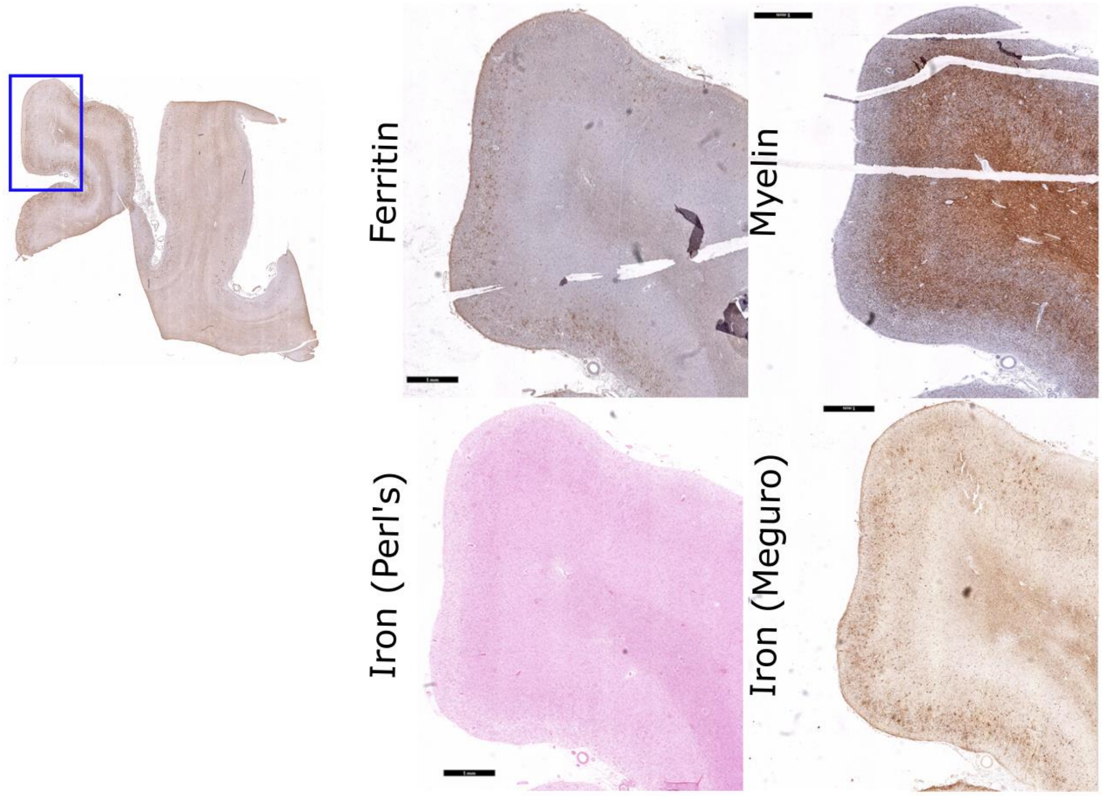


***Figure S2*. *Histopathology on the AD tissue block.*** *Close-up on the gyrus. Scalebar on the close-up is 1 mm.*

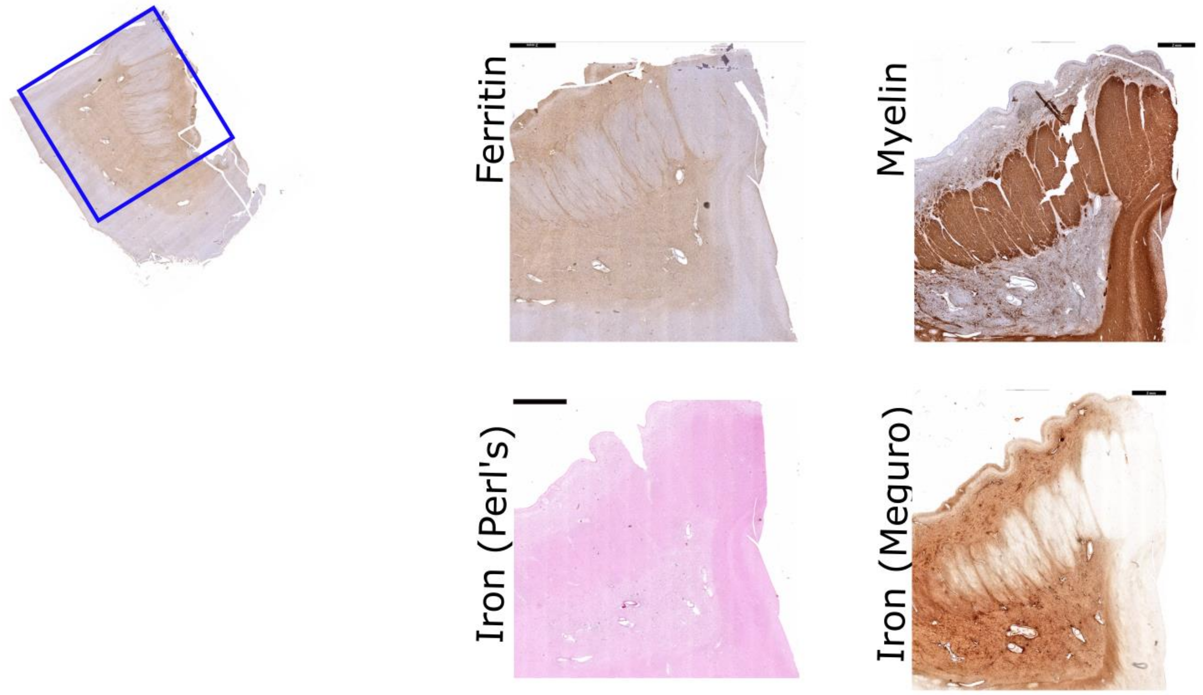


***Figure S3*. *Histopathology on the HD tissue block.*** *Close-up on putamen and caudate nucleus. Scalebar on the close-up is 1 mm.*

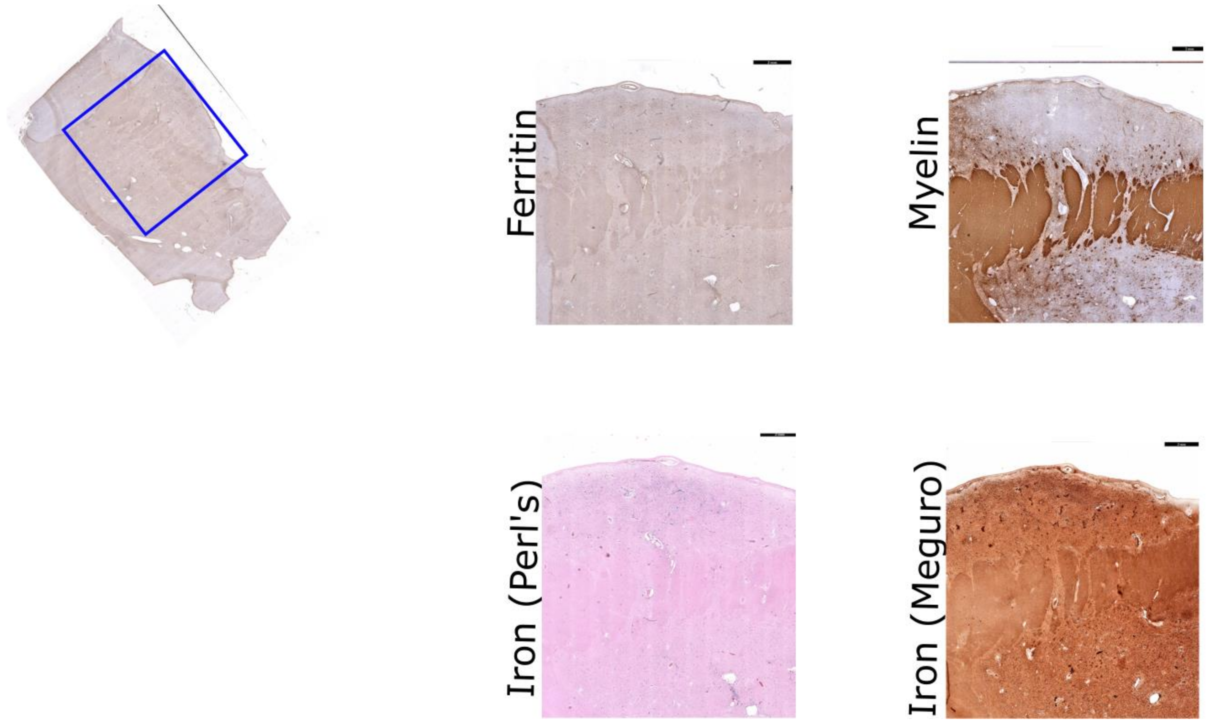


***Figure S4. Histopathology on the ACP tissue block.*** *Close-up on the internal capsule, caudate nucleus and part of the putamen. Scalebar on the close-up is 1 mm.*

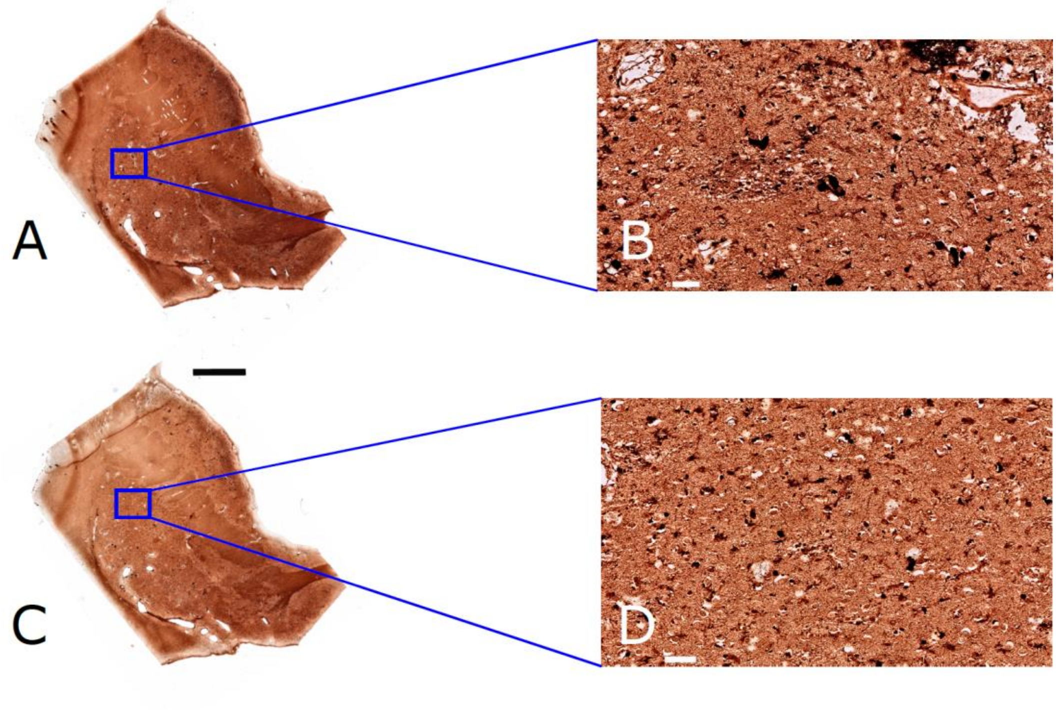


***Figure S5. Illustration of the iron accumulation in aceruloplasminemia brain.*** *Panels A and C show the modified Meguro staining on the whole tissue slice. Panel A was obtained on a 20 µm slice and a protocol optimized for the slice thickness. Panel C was obtained on an adjacent slice of 20 µm, but with shorter incubation times (the protocol was optimized for a 5 µm protocol slides). Panel B and D are close-ups on the region of interest in the putamen (blue square). The images illustrate the exceptional degree of iron accumulation in ACP. Scale bar is 50 µm in B and D, and 5 mm in A and C.*
